# BLISS: quantitative and versatile genome-wide profiling of DNA breaks *in situ*

**DOI:** 10.1101/091629

**Authors:** Winston X. Yan, Reza Mirzazadeh, Silvano Garnerone, David Scott, Martin W. Schneider, Tomasz Kallas, Joaquin Custodio, Erik Wernersson, Linyi Gao, Yinqing Li, Yana Federova, Bernd Zetsche, Feng Zhang, Magda Bienko, Nicola Crosetto

**Author notes:** Equally contributing authors (Winston X. Yan, Reza Mirzazadeh). Corresponding authors: N.C., M.B., and F.Z.

## Abstract

We present a method for genome-wide DNA double-strand Breaks (DSBs) Labeling *In Situ* and Sequencing (BLISS) which, compared to existing methods, introduces several key features: 1) high efficiency and low input requirement by *in situ* DSB labeling in cells or tissue sections directly on a solid surface; 2) easy scalability by performing *in situ* reactions in multi-well plates; 3) high sensitivity by linearly amplifying tagged DSBs using *in vitro* transcription; and 4) accurate DSB quantification and control of PCR biases by using unique molecular identifiers. We demonstrate the ability to use BLISS to quantify natural and drug-induced DSBs in low-input samples of cancer cells, primary mouse embryonic stem cells, and mouse liver tissue sections. Finally, we applied BLISS to compare the specificity of CRISPR-associated RNA-guided endonucleases Cas9 and Cpf1, and found that Cpf1 has higher specificity than Cas9. These results establish BLISS as a versatile, sensitive, and efficient method for genome-wide DSB mapping in many applications.

---

DNA double-strand breaks (DSBs) are major DNA lesions that form in a variety of physiological conditions – such as transcription^1
,2^, meiosis^3^ and VDJ recombination^4^ – and arise as a consequence of DNA damaging agents and replication stress^5^. DSBs can also be induced in a controlled fashion at specific sites in the genome using programmable nucleases such as CRISPR (clustered regularly interspaced short palindromic repeats)-associated RNA-guided endonucleases Cas9 and Cpf1, which have greatly advanced genome editing. However, the potentially mutagenic off-target DNA cleavage activities of these nucleases represent a major concern that needs to be evaluated before these enzymes can be used in the clinical setting^6^. Thus, developing methods that can accurately map the genome-wide location of endogenous, as well as induced DSBs in different systems and conditions is not only essential to expand our understanding of DSB biology, but is also crucial to enable the translation of programmable nucleases into clinical applications.

In the past few years, several methods based on next-generation sequencing have been developed to assess DSBs at genomic scale, including ChIP-seq^7,8^, BLESS^9–11^, GUIDEseq^12^, Digenome-seq^13^, IDLV-mediated DNA break capture^14^, HTGTS^15^, and more recently End-Seq^16^ and DSBCapture^17^. While in general all of these methods represent complementary tools (**Supplementary Table 1**), they also have important drawbacks. For example, ChIP-seq for factors such as γH2A.X cannot label DSBs directly or identify breakpoints with single-nucleotide resolution. GUIDEseq, IDLV-mediated DNA break capture, and HTGTS detect DSBs by quantifying the products of non-homologous end-joining (NHEJ) repair, potentially missing DSBs repaired through other pathways. Furthermore, *in vivo* delivery of exogenous oligos in GUIDEseq or viral cassettes in IDLV-mediated DNA break capture is challenging, especially for primary cells and intact tissues. DSBs induced by programmable nucleases can be evaluated *in vitro* using Digenome-seq, but this approach may not be representative of physiologically relevant conditions – such as chromatin environment and nuclear architecture – in controlling the frequency of DNA breaking and repair, or of relevant nuclease concentrations. Lastly, BLESS and its recent modifications, End-Seq^16^ and DSBCapture^17^, require large quantities of input material, are labor-intensive, and are not quantitative due to lack appropriate controls for amplification biases, which altogether limits their applications and scalability.

We present here a more versatile, sensitive and quantitative method for detecting DSBs applicable to low-input specimens of both cells and tissues that is scalable for high-throughput DSB mapping in multiple samples. Our method, Breaks Labeling *In Situ* and Sequencing (BLISS), features efficient *in situ* DSB labeling in fixed cells or tissue sections immobilized onto a solid surface, linear amplification of tagged DSBs via T7-mediated *in vitro* transcription (IVT)^18^ for greater sensitivity, and accurate DSB quantification by incorporation of unique molecular identifiers (UMIs)^19^ (Fig. 1a, Supplementary Fig. 1a, Supplementary Table 2, Online Methods and Supplementary Information).

**Figure 1.**
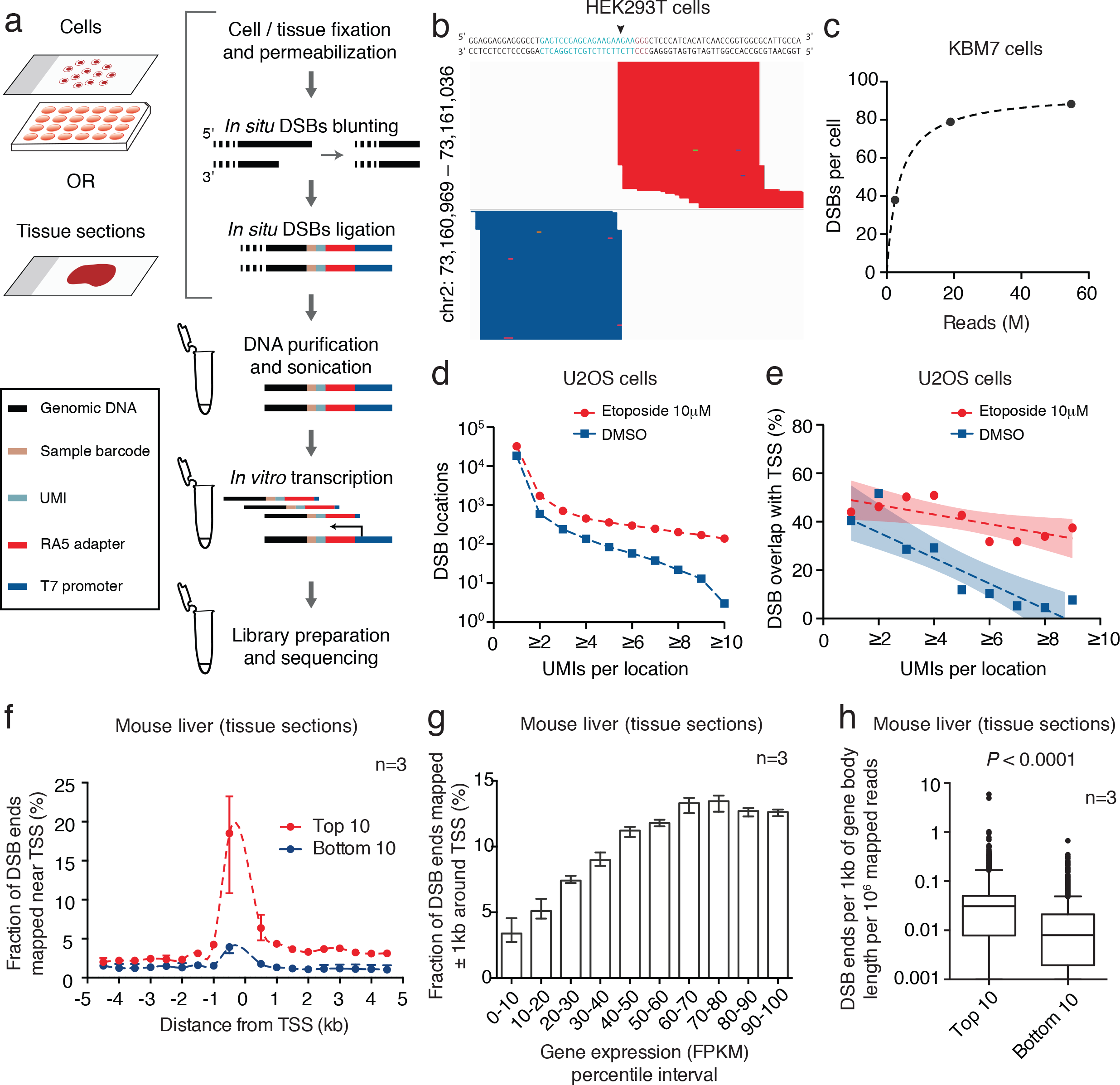
Quantitative detection of natural and drug-induced double-strand breaks (DSBs). (**a**) Schematic of BLISS. The workflow starts by either fixing cells onto a microscope slide or in a multi-well plate, or by immobilizing already fixed tissue sections onto a slide. DSB ends are then *in situ* blunted and tagged with dsDNA adapters containing components described in the boxed legend and in **Supplementary Table 2**. Tagged DSB ends are linearly amplified using *in vitro* transcription, and the resulting RNA is used for Illumina library preparation and sequencing. (**b**) BLISS reads aligned to an SpCas9 on-target cut site (arrowhead) in the *EMX1* gene. Light blue, guide sequence. Orange, PAM sequence. Dark blue, reads mapped to the minus strand. Red, reads mapped to the plus strand. (**c**) Estimated number of DSBs per cell in three replicates sequenced at increasing sequencing depth. Dashed line, hyperbolic interpolation. (**d**) Number of DSB locations in etoposide-treated versus control U2OS cells by filtering on the minimum number of unique molecular identifiers (UMIs) per DSB location. (**e**) Fraction of DSB locations mapped around the transcription start sites (TSS) in control versus etoposide-treated U2OS cells as a function of the minimum number of UMIs per DSB location. Dashed lines, linear interpolation. Color shades, 95% confidence intervals. (f) For BLISS on mouse liver, mapping of sequenced DSB ends found in the top 10% (red) and bottom 10% (blue) of expressed genes in the mouse liver. n, number of biological replicates. Dots, mean value. Whiskers, min-max range. Dashed lines, spline interpolation. (g) Percentage of sequenced DSB ends mapped in a ±1kb interval around the TSS for each inter-decile interval of gene expression in mouse liver. FPKM, fragments per kilobase of transcript per million mapped reads. n, number of biological replicates. Bars, mean value. Whiskers, min-max range. (**h**) Number of sequenced DSB ends mapped per kilobase inside the gene body of the top 10% and bottom 10% expressed genes in mouse liver. n, number of biological replicates. Whiskers, 2.5–97.5 percentile range. *P*, Mann-Whitney test.

We performed multiple BLISS experiments in various sample types and preparations, including low-input samples of cells and tissue sections, drug treatments, and nuclease treatments with CRISPR-Cas9 and CRISPR-Cpf1, obtaining high-quality libraries with balanced UMI and strand composition (**Supplementary Fig. 1b-d and Supplementary Table 3**). We developed a robust sequencing data processing pipeline that, by using the information contained in the UMIs, enables us to reliably distinguish DSB events that have occurred at the same genomic location in different cells (**Supplementary Fig. 1e-g and Supplementary Information**).

To test whether BLISS can faithfully detect DSBs in the genome, we first transfected HEK293T cells with *Streptococcus pyogenes* Cas9 (SpCas9) and a single-guide RNA (sgRNA) targeting the *EMX1* gene, for which BLISS was able to precisely localize and quantify both DSB ends generated by SpCas9 at the correct on-target location (Fig. 1b). Furthermore, in low-input samples of KBM7 cells, BLISS precisely identified telomeric ends – which mimic DSB ends – confirming our previous results using BLESS in a much larger number of cells^9^ (**Supplementary Fig. 2a**).

**Figure 2.**
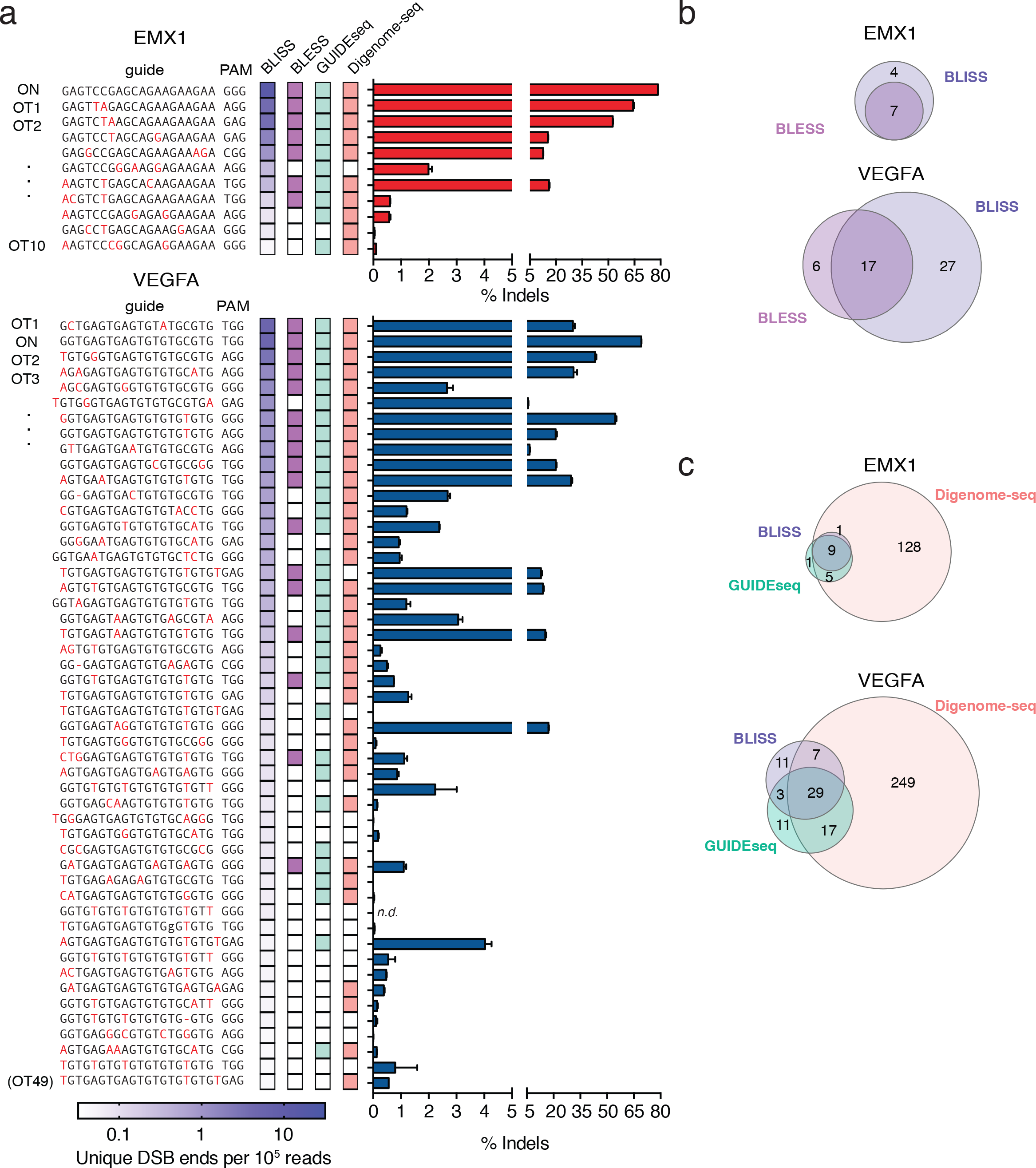
Genome-wide quantification of SpCas9 on- and off-target DSBs. (**a**) On- and off-target sites identified by BLISS, BLESS, GUIDEseq, and Digenome-seq. BLISS targets were ranked in descending order based on the number of unique DSB ends aligned to the target per 10^5^ unique BLISS reads. Colors in the BLESS, GUIDEseq, and Digenome-seq columns indicate when the BLISS target was previously found by either of these methods. Individual sites were validated by targeted deep sequencing and the percentage of reads containing an insertion or deletion (indel) is shown. (n=3, error bars show S.E.M.). ON, on-target. OT, off-target. (**b**) Overlap between on- and off-target sites identified by BLISS versus BLESS. (**c**) Overlap between on- and off-target sites identified by BLISS versus GUIDEseq and Digenome-seq.

We then assessed the accuracy of BLISS by sequencing at increasing depth three libraries obtained from low-input samples of KBM7 cells (**Supplementary Table 3**). By performing rarefaction analysis on the number of unique DSBs labeled by UMIs detected at increasing sequencing depths, we estimated that BLISS was able to detect 80-100 DSBs per cell (Fig. 1c and Online Methods). This estimate was within the same range of the number of γH2A.X foci quantified by microscopy in the same cell line (85.7 ± 60.6 foci/cell, mean ± s.d., **Supplementary Fig. 2b and 2c**), suggesting that most of the DSBs detected by BLISS represent true biological events rather than background noise.

To further assess the quantitative nature of BLISS, we used UMIs to quantify DSBs induced by the topoisomerase inhibitor, etoposide. In two biological replicates of U2OS cells treated with etoposide, the number of unique DSB ends detected by BLISS increased in a dose-dependent manner, consistent with γH2A.X measurements (**Supplementary Fig. 3a-c**). The treatment resulted in DSB accumulation at recurrent genomic locations in multiple cells, which could be distinguished because multiple DSB ends mapping to the same location were labeled by distinct UMIs (Fig. 1d and Supplementary Fig. 3d and 3e). These recurrent locations were significantly enriched in the neighborhood of transcriptional start sites (TSS), confirming prior findings by BLESS that etoposide has prominent effects around TSS^20^ (Fig. 1e **and Supplementary Fig. 3f**).

**Figure 3.**
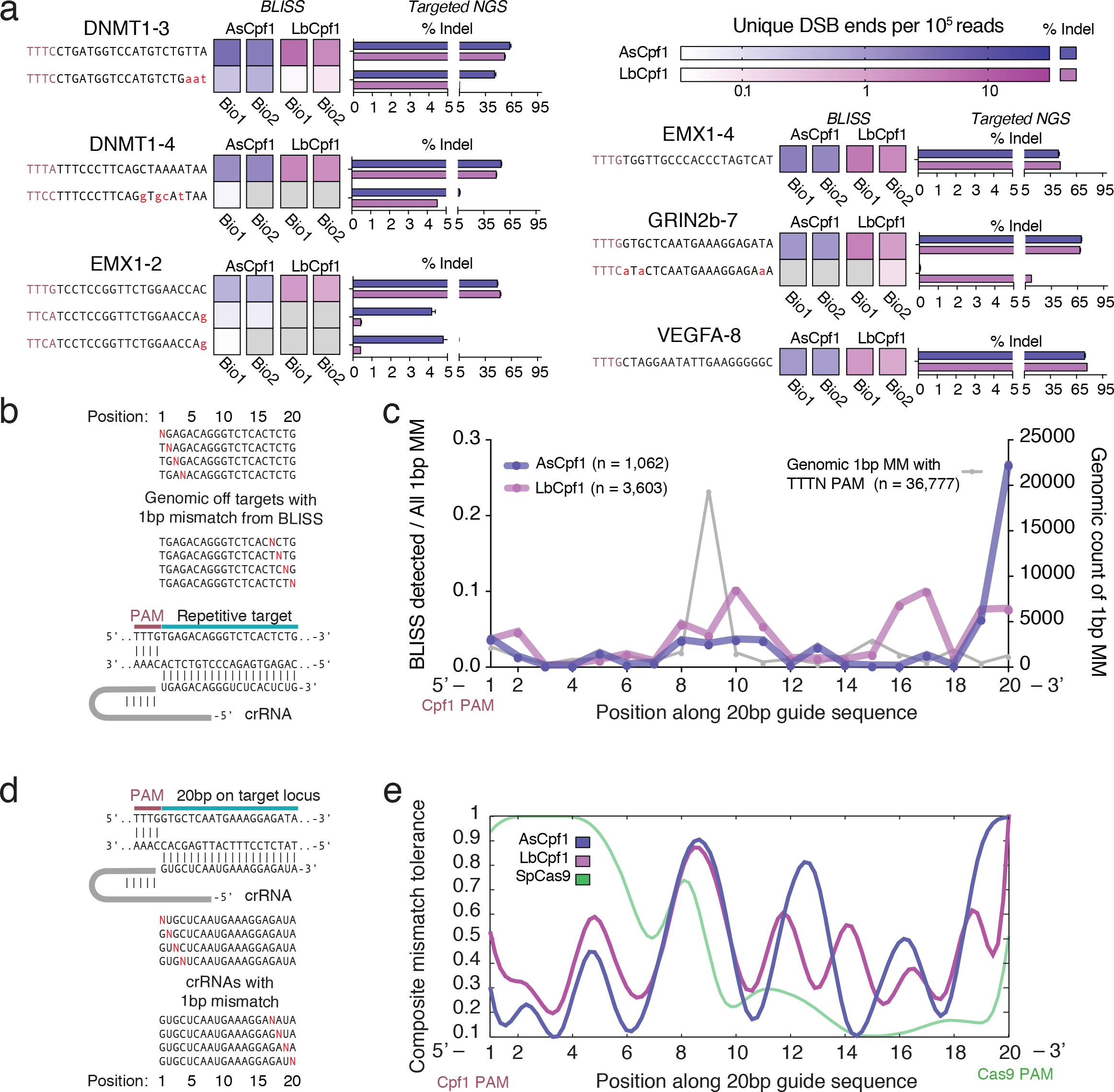
Characterization of AsCpf1 and LbCpf1 specificity. (**a**) Validated on- and off-target sites for AsCpf1 and LbCpf1 for six separate guide targets as measured by Cpf1-BLISS over two independent biological replicates and validated by targeted NGS (*n=3*, error bars show S.E.M.). Gray boxes indicate DSB loci not detected within a biological replicate. (**b**) Evaluating the position-dependent mismatch tolerance of AsCpf1 and LbCpf1 using a repetitive guide with 36,777 predicted genomic loci with single mismatches. (**c**) A map of mismatch tolerance per position generated by dividing at each base the number of off-targets discovered in BLISS versus the possible single mismatched genomic targets for Cpf1. The gray line plotted on the left y-axis is the count of single mismatched targets in the genome for Cpf1 as predicted by Cas OFFinder^22^. (**d**) Guide designs for investigating the effect of single base pair mismatches in the RNA guide on AsCpf1 and LbCpf1 specificity by measuring the change in their on-target efficiency versus a matched guide. (**e**) Composite mismatch tolerance model for AsCpf1 and LbCpf1 based on saturated single base pair mismatches for two guides. Cas9 data (green) modeled from existing Cas9 single mismatch data^25^.

The ability to obtain genome-wide DSB maps from primary cells and tissue samples would greatly help studies of DNA damage and repair processes in animal models and clinical samples. With this goal in mind, we performed proof-of-principle experiments using either tissue sections or nuclei derived from mouse liver biopsies (**Supplementary Fig. 4a and 4b**). In line with recent findings in different cell types^1,2,17^, DSBs were strongly enriched in the neighborhood of TSS as well as along the gene body of highly expressed genes (Fig. 1f-h and Supplementary Fig. 4c-e). Gene ontology analysis of those genes that were reproducibly identified as carrying the highest DSB levels in three biological replicates revealed a significant enrichment in functional terms related to liver-specific metabolic processes, indicating that BLISS is able to capture endogenous DSBs related to tissue-specific processes (**Supplementary Fig. 4f and Supplementary Tables 4 and 5**). Similar findings were recapitulated in low-input samples of primary mouse embryonic stem cells (**Supplementary Fig. 4g-i**), confirming that BLISS is a highly versatile method that can be applied to study endogenous DSBs in a range of cell and tissue samples. Furthermore, we assessed chromatin accessibility in the same tissue samples by applying a modified BLISS protocol in which DNA breaks were introduced *in situ* by the HindIII restriction endonuclease (**Supplementary Fig. 5a and 5b and Online methods**). This revealed that, even though DSBs tend to form more in open chromatin as previously reported^16,17^, genomic regions with similar chromatin accessibility can have very different DSB levels and *vice versa* (**Supplementary Fig. 5c**).

We next aimed to assess the sensitivity of BLISS by characterizing the DSBs induced by the CRISPR-associated RNA-guided endonucleases, Cas9 and Cpf1. Evaluating Cas9 and Cpf1 on and off-targets is a valuable way of assessing BLISS sensitivity because the nuclease-induced cleavage sites 1) are sparse enough so as to not saturate BLISS; 2) are relatively well-defined by both cut site location determined by other assays as well as homology to the original target; and 3) occur over a wide dynamic range of DSB frequencies to allow quantification of the detection sensitivity. Meanwhile, BLISS is a versatile and minimally disruptive technique for studying the specificity of CRISPR nucleases since by labeling DSBs post fixation, it requires no additional perturbations to the cell beyond delivery of the nuclease and RNA guide. Hence, we developed a workflow to screen the off-target activity of Cas9 or Cpf1 endonucleases using BLISS (Cas9-BLISS/Cpf1-BLISS) in parallel with existing genome editing protocols (**Supplementary Fig. 6a**). Aside from culturing cells for BLISS on poly-D-lysine coated plates and fixation 24 hours post transfection, no additional modifications of delivery reagents or workflows were necessary, allowing BLISS to capture a snapshot of the CRISPR activity in cells with minimal bias.

To benchmark the sensitivity of Cas9-BLISS against existing genome-wide specificity methods such as BLESS, GUIDEseq, and Digenome-seq, we transfected HEK293T cells with SpCas9 and two sgRNAs targeting the *EMX1* and *VEGFA* genes, both of which have been characterized using all three methods^11,12,14,15^. This set of known off-targets allowed us to further optimize Cas9-BLISS through direct comparison of different DSB labeling strategies, showing that *in situ* A-tailing prior to adapter ligation increased the sensitivity of DSB detection when directly compared to the original blunt end ligation chemistry (**Supplementary Fig. 6b-e**). We furthermore refined the computational pipeline that we previously established for identifying *bona fide* Cas9 DSBs for the analysis of Cas9-BLESS data^10^ to achieve greater sensitivity (**Online Methods**). In addition to the expected on-target DSB sites, BLISS detected numerous off-target sites that were successfully validated by targeted next-generation sequencing (NGS), including many sites previously identified by BLESS, GUIDEseq, or Digenome-seq (Fig. 2a and Supplementary Table 6). BLISS also uncovered new off-target sites that were not found when the refined computational pipeline was re-applied to published BLESS data on the same targets^11^ (Fig. 2b). Side-by-side comparison of BLISS with Digenome-seq and GUIDEseq revealed that while all three methods generally agree on the top off-targets identified, they differ in the number of weaker off-target sites, particularly in the case of *VEGFA* (Fig. 2c).

We next applied BLISS to characterize the DNA-targeting specificity of the recently described CRISPR-associated endonuclease Cpf1 (Cpf1-BLISS). Cpf1 is a two-component RNA-programmable DNA nuclease with several unique properties that may broaden the applications of genome engineering: 1) it employs a short CRISPR RNA (crRNA) without an additional trans-activating CRISPR RNA (tracrRNA); 2) it utilizes a T-rich protospacer-adjacent motif (PAM) located 5’ to the target sequence; and 3) it additionally generates a staggered cut with a 5’ overhang^21^. We selected six Cpf1 targets across four different genes for genome-wide off-target evaluation using BLISS and targeted NGS. Four targets have NGG PAMs on the 3’ end to enable a simultaneous comparison between SpCas9 and eSpCas9. We evaluated Cpf1 from *Acidaminococcus sp.* (AsCpf1) and *Lachnospiraceae bacterium* (LbCpf1), both which have been harnessed for efficient mammalian genome editing^21^.

At the dual Cpf1 and Cas9 targeted loci, BLISS revealed differences in the *in vivo* pattern of DSBs induced by these two enzymes. Taking the histogram of all the differences between reads mapping to opposite sides of DSBs (**Supplementary Fig. 7a**) showed that while Cas9 cuts are generally blunt ended or contain 1nt overhangs, Cpf1 cuts exhibit a wide distribution of overhang lengths depending on the target (**Supplementary Fig. 7b**). Since *in vitro* cleavage of AsCpf1 and LbCpf1 reveals 4–5nt 5’ overhangs as the predominant cleavage outcome^21^, these results suggest that *in vivo* processing of Cpf1 cut sites generates heterogeneous DSB patterns.

To identify Cpf1 off-target sites using BLISS, we applied the same computational pipeline as was used for Cas9-BLISS. We performed targeted NGS on all off-target sites identified from independent BLISS biological replicates from both AsCpf1 and LbCpf1 to maximize sensitivity (**Supplementary Fig. 8**). Comparing the BLISS results for AsCpf1 or LbCpf1 with SpCas9, we consistently found fewer *bona fide* off-target sites for the two Cpf1 orthologs (Fig. 3a and Supplementary Fig. 8), suggesting that Cpf1 is less tolerant of mismatches than Cas9.

For the four targets with shared Cpf1 and Cas9 PAMs, genome modification with SpCas9 yielded a greater range of *bona fide* off-target sites (**Supplementary Fig. 9**), consistent with prior observations that individual SpCas9 guides can have a wide variation in the number of off-target sites independent of the prevalence of closely matched sites in the genome^12^. As expected, the use of eSpCas911 reduced the number of off-targets without loss of on-target activity.

Lastly, to assess whether BLISS was sensitive enough to detect a large number of Cpf1-induced breaks across a wide dynamic range of cleavage activity, we designed additional guides for Cpf1, targeting repetitive sequences with 278 (*GRIN2b* repetitive guide) and 8,130 (*DNMT1* repetitive guide) perfectly matched on-target sites with a TTTN PAM, as predicted using Cas-OFFinder^22^. A wide range of both on- and off-target loci were detected using Cpf1-BLISS (**Supplementary Fig. 10**), suggesting that Cpf1 can indeed have a high level of specificity for guides not targeting repetitive regions. Altogether, these results corroborate the findings of other recent studies that Cpf1 can be highly specific_23,24_.

The Cpf1 repetitive targets also enabled us to study the position dependence of mismatch tolerance by examining whether mismatches in certain positions are enriched in the off-target results versus the genomic background. In particular, the *DNMT1* repetitive guide has nearly 37,000 off-targets with a single mismatch to the on-target sequence and a TTTN PAM, according to Cas-OFFinder^22^. Each mismatched position is represented in at least 150 genomic loci, though the prevalence of a mismatch at a given target position is not uniformly distributed (Fig. 3b-c). Cpf1-BLISS detected ~1,000 and ~3,600 off-targets for AsCpf1 and LbCpf1, respectively, that contain only one mismatch to the on-target sequence. The fraction of Cpf1-BLISS-detected sites over all possible mismatches at that position was calculated to obtain a measure of how permissive Cpf1 is to mismatches along the guide (Fig. 3c). We also systematically introduced mismatches between the Cpf1 guide and target DNA, normalizing the on-target modification rate for each mismatched guide to the matched target (Fig. 3d and Supplementary Fig. 11). The on-target indel data from the mismatched guides were used to generate a composite model of the mismatch tolerance versus position for AsCpf1 and LbCpf1 (Fig. 3e and Supplementary Methods), with the overlaid SpCas9 trace based on reanalysis of previous mismatch data^25^ (**Supplementary Methods**). Taken together, there appears to be three regions of the guide for both AsCpf1 and LbCpf1 where mismatches are more tolerated: 1) at the 3’ PAM distal end of the guide (positions 19-20); 2) towards the middle of the guide (positions 8-11); and, to a lesser degree, 3) at the first base at the 5’ PAM proximal end (position 1). This qualitatively suggests that Cpf1 may have several distinct regions of the guide that enforce complementarity and thereby contribute to its heightened specificity compared to SpCas9.

In conclusion, we have developed a sensitive method for direct genome-wide DSB detection that compared to the existing methods presents major improvements: 1) robust quantification by using UMIs to reduce data noise and count DSBs that form in different cells at the same genomic location; 2) applicability to low-input cell and tissue samples by performing all *in situ* reactions and washes on a solid surface (if sample input is not limiting, *in situ* reactions can also be performed in-tube, thus providing additional versatility); 3) assay scalability and cost-effective multiplexing by performing *in situ* reactions inside multi-well plates and barcoding samples in different wells before pooling; 4) fast turnaround time compared to BLESS (~12 active work-hours over 5 days to process 24 samples by BLISS versus at least ~60 active work-hours over 15 days by BLESS). Finally, we demonstrate that BLISS is a highly sensitive method to assess the DNA-targeting specificity of CRISPR-associated RNA-guided DNA endonucleases Cas9 and Cpf1, and we show that, in agreement with previous reports^23,24^, Cpf1 can provide high levels of editing specificity. These features make BLISS a powerful and versatile method for genome-wide DSB profiling that we believe will advance the study of natural and artificially induced DSBs in many conditions and model systems, including precious clinical samples.

## ACKNOWLEDGEMENTS

W.X.Y. is supported by T32GM007753 from the National Institute of General Medical Sciences and a Paul and Daisy Soros Fellowship. E.W. is a recipient of a Swedish Society for Medical Research (SSMF) Postdoctoral Fellowship. F.Z. is supported by the National Institutes of Health (5DP1-MH100706, 1RO1-MH110049-02, 1 R21 EY026412-01), a Waterman Award from the National Science Foundation, Howard Hughes Medical Institute, the New York Stem Cell, Simons, Paul G. Allen Family, and Vallee Foundations, and James and Patricia Poitras, Robert Metcalfe, and David Cheng. F.Z. is a New York Stem Cell Foundation Robertson Investigator. M.B. is supported by the Science for Life Laboratory, the Swedish Research Council (621-2014-5503), and the Ragnar Söderberg Foundation. N.C. is supported by the Karolinska Institutet, the Swedish Research Council (521-2014-2866), the Swedish Cancer Research Foundation (CAN 2015/585), and the Ragnar Söderberg Foundation.

## EXPERIMENTAL METHODS

### Cell and tissue samples

The following cell lines were used: KBM7 from Oscar Fernandez-Capetillo (SciLifeLab, Stockholm, Sweden); U2OS from Mats Nilsson (SciLifeLab, Stockholm, Sweden); HEK 293T from ATCC; mouse embryonic stem cells (mESCs) from Simon Elsaesser (SciLifeLab, Stockholm, Sweden). Culturing conditions were as following: KBM7 in Iscove’s modified Dulbecco’s medium (IMDM, Life Technologies, cat. no. 10829018), supplemented with 10% fetal bovine serum (FBS, Gibco, cat. no. F2442); U2OS in Dulbecco’s modified Eagle’s medium (DMEM, Life Technologies, cat. no. D0819), supplemented with 10% FBS; HEK293T in DMEM supplemented with 10% FBS; mESCs in minimal essential medium (MEM, Sigma, cat. no. M2279), supplemented with 20% FBS, 1% GlutaMAX (Gibco, cat. no. 35050061), 1% nonessential amino acids (Gibco, cat. no. 11140035), 1% sodium pyruvate (Gibco, cat. no. 11360070), and 0.2% β-mercaptoethanol, in the presence of leukemia inhibitory factor (Sigma cat. no. L5158-5UG) corresponding to 1,000 U/ml. All cell lines were tested to be mycoplasma free using MycoAlert™ Mycoplasma Detection Kit (Lonza, cat. no. LT07-118). For tissue-BLISS on mouse liver, wild-type 6-weeks old C57/BL6 male mice were sacrificed following the guidelines in the MIT protocol #0414-027-17 ‘Modeling and Treating Genetic Disease Using Targeted Genome Engineering’ (IACUC AWA #A3125-01, IACUC #0411-040-14, approval date 5/16/2013).

### Cas9/Cpf1 expression constructs and transfections

The selected targets for Cas9-BLISS are located within the EMX1 locus (GAGTCCGAGCAGAAGAAGAA gGG) and the VEGFA gene locus (GGTGAGTGAGTGTGTGCGTG tGG). The plasmids used containing the SpCas9 and the sgRNA cassette were identical to the ones used for Cas9-BLESS11, where the targets were labeled as EMX1(1) and VEGFA(1). The same targets have also been studied using GUIDEseq^12^, where they were labeled as EMX1 and VEGFA_site3. AsCpf1 and LbCpf1 along with their cognate crRNAs were cloned into the same expression vector as Cas9 to enable a direct comparison. Cells were plated before transfection in 24-well plates pre-coated with poly-D-lysine (Merck Millipore, cat. no. A003E) at a density of ~125,000/well, and were let grow for 16-18h until 60–70% confluence. For transfections, we used 2μl of Lipofectamine 2000 (Life Technologies, cat. no. 11668019) and 500ng of Cas9 plasmid in 100μl total of OptiMEM (Gibco, cat. no. 31985062) per each well of a 24-well plate as described previously (Ran et al., 2015).

### γH2A.X immunofluorescence

γH2A.X immunostaining was performed as previously described^9^, using a mouse anti-phospho-histone H2A.X (ser139) primary antibody (Millipore, cat. no. 05-636) diluted 1:1000 in blocking buffer and a goat anti-mouse IgG (H+L) Alexa Fluor^®^ 647 conjugate (Thermo, cat. no. A-21235) secondary antibody diluted 1:1000 in blocking buffer. To image γH2A.X foci, we acquired images every 0.4μm throughout the entire nuclear volume using a 40× oil objective and a LSM 780 confocal microscope (Zeiss).

### BLISS adapters

All BLISS adapters were prepared by annealing two complementary oligonucleotides as described below. All oligos were purchased from Integrated DNA Technologies as standard desalted oligos. UMIs were generated by random incorporation of the four standard dNTPs using the “Machine mixing” option. Before annealing, sense oligos diluted at 10μM in nuclease-free water were phosphorylated for 1h at 37°C with 0.2U/μl of T4 Polynucleotide Kinase (NEB, cat. no. M0201). Phosphorylated sense oligos were annealed with the corresponding antisense oligos pre-diluted at 10μM in nuclease-free water, by incubating them for 5min at 95°C, followed by gradual cooling down to 25°C over a period of 45min (1.55°C/min) in a PCR thermo-cycler.

### BLISS

A step-by-step BLISS protocol is provided in **Supplementary Information**. For BLISS in cell lines, we typically either grew cells directly onto 13mm coverslips (VWR, cat. no. 631-0148) or we spotted them onto coverslips pre-coated with poly-L-lysine (PLL) (Sigma, cat. no. P8920-100ML). For Cas9 & Cpf1 experiments, we fixed HEK293T cells directly into the 24-well plate used for transfections, and performed all *in situ* reactions done directly inside the plate. For BLISS in mouse liver, we developed two approaches: 1) Tissue cryopreservation and sectioning. Freshly extracted liver biopsies were first fixed in PFA 4% for 1h at rt, and then immersed in a sucrose solution (15% overnight and then 30% until the tissue sank) before embedding in OCT. 30μm-thick tissue sections were mounted onto microscope slides, dried for 60min at rt, and stored at 4°C before further processing. 2) Preparation of nuclei suspensions. Freshly extracted liver biopsies were cut into small pieces and transferred into a 1.5-2ml tube containing nucleus isolation buffer (NaCl 146mM, Tris-HCl 10mM, CaCl_2_ 1mM, MgCl_2_ 21mM, Bovine Serum Albumin 0.05%, Nonidet P-40 0.2%, pH 7.8). We typically incubated the samples for 15-40min until the tissue fragments became transparent, after which the nuclei were centrifuged for 5min at 500g and then re-suspended in 200-500μl of 1*×* PBS. 100μl of nuclei suspension were dispensed onto a 13mm PLL-coated coverslip and incubated for 10min at rt. Afterwards, 100μl of PFA 8% in 1*×* PBS were gently added and incubated for 10min at rt, followed by two washes in 1*×* PBS at rt. The samples were stored in 1*×* PBS at 4°C up to one month before performing BLISS.

### *In situ* DNA digestion

Samples for DNA accessibility mapping were prepared in the same way as BLISS samples, except that the *in situ* DSBs blunting step was substituted by an *in situ* DNA digestion step using 1U/μl of HindIII endonuclease (NEB, cat. no. R3104) and incubating the samples for 18h at 37°C. HindIII cut sites were ligated with modified BLISS adapters carrying the HindIII complementary sticky end (see **Supplementary Fig. 4h**). In order to prevent *in situ* re-ligation of HindIII cut sites, the samples were incubated for 2h at 37°C in the presence of 0.015U/μl of Calf Intestinal Alkaline Phosphatase (Promega, cat. no. M2825) before *in situ* ligation.

## COMPUTATIONAL METHODS

### Image processing and counting of γH2AX foci and cells

All algorithms were implemented in MATLAB using custom-made scripts, available upon request. To count γH2AX foci in KBM7 cells, we first segmented nuclei stained with DAPI using image thresholding. We then identified all local maxima within each image and then ranked the maxima according to their response to a Laplacian filter. We then fitted a Gaussian to the first peak of the histogram of the filter responses, corresponding to background noise (i.e., autofluorescence and photon noise). We counted γH2AX foci per nuclei using the dots with a filter response of more than 10 standard deviations above the mean of the background. To count cells prior to capture and genomic DNA extraction, we first rinsed samples in nuclease-free water, air-dried them, and acquired wide-field images of areas selected for cell capture using a TI-S-E Motorized stage operated by NIS-Elements software (Nikon). Next, we identified objects in wide-field images by locating maxima of the determinant of the gradient structure tensor. We then classified objects being cells or not based on anisotropy, size, and median gradient magnitude. Finally, we manually corrected and verified the segmentation.

### Pre-processing of sequencing data

To convert the raw sequencing data into BED files ready to be used for *ad hoc* analyses, we applied the pipeline summarized in **Supplementary Fig. 1e**. Briefly, we filtered the FASTQ files for overall quality by requiring a Phred score ≥30 for every base. Thereafter, we scanned the filtered reads for the presence of the exact prefix (8N UMI and sample barcode), by allowing up to 2 mismatches in the UMI portion and up to 1 mismatch in the barcode (see the analysis of UMI errors in **Supplementary Information**). After removal of the prefix, we aligned the reads to the reference genome (GRCh37/hg19 for human, NCBI37/mm9 for mouse). We retained reads mapping with a quality score ≥5, after excluding regions with poor mappability (https://www.encodeproject.org/annotations/ENCSR636HFF/). Next, we performed a further filtering step based on UMI sequences to filter out PCR duplicates. Reads mapping in nearby locations (at most 8nt apart) and having at most 2 mismatches in the UMI sequence were associated with the location of the most frequent read in the neighborhood. Finally, we generated BED files containing a list of genomic locations associated with unique UMIs to be used in downstream analyses.

### Identification of telomeric ends among sequenced reads

To analyze the composition of BLISS reads derived from the telomeric C-rich strand, we screened R1 reads with the correct prefix (8N UMI and sample barcode) for the presence of each of the 6 possible patterns based on the human telomeric sequence: [#A,#AA,#TAA,#CTAA,#CCTAA,#CCCTAA]-CCCTAA.

### Estimation of DSBs per cell

To estimate the number of spontaneous DSBs, we sequenced at different depth three libraries prepared from small numbers of cultured KBM7 cells (L1, L3 and L4, see **Supplementary Table 3**). For each sample, we estimated the number of DSBs per cell by counting the number of sequenced reads with correct prefix mapped to a unique genomic location and tagged by a unique UMI, and assuming that on average one DSB produces 2 unique reads. We then fitted the data to the model 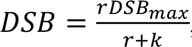, where *DSB_max_* is the number of DSB events per cell at saturation, *r* is the number of total reads, and *k* is a constant. At saturation, the model estimated *DSBmax*=94 breaks per cell (95% C.I. 93.10–95.07), in agreement with γH2A.X foci counting in the same cell line (**Supplementary Fig. 2c**).

### Quantification of etoposide effects

For U2OS cells treated with etoposide, we counted the number *n* of unique DSB locations on each chromosome that were found with at least 1≤*t*≤10 UMIs and at most t=500 UMIs. We then normalized the cumulative sum, *n* by the total number of DSB ends sequenced, and calculated the ratio between the normalized cumulative sum in the treated and non-treated sample, and averaged the fold change over all chromosomes. We repeated the same process separately for the unique locations exclusively found in the etoposide-treated or untreated sample. For enrichment analysis of etoposide-induced DSBs around the TSS, we calculated the fraction of unique DSB locations (found with at least 1≤*t*≤10 UMIs and at most t=500 UMIs) that fell in a window of ±5kb centered on the TSS of all genes.

### Quantification of spontaneous DSBs near TSS and within gene bodies

For mouse liver and mESCs, we used RNA-seq data obtained from the Mouse Encode Project at Ren lab (http://chromosome.sdsc.edu/mouse/download.html. We first identified the top 10% and bottom 10% expressed genes and then, for each gene in the two groups, we calculated the number of unique DSB ends (i.e., the number of DSB locations on either strand associated with a unique UMI) falling in an interval of ±5kb centered on the TSS of the gene. This approach enabled us to distinguish DSBs that had occurred at the same genomic location in different cells. We then calculated the proportion of all the DSB locations mapped around the TSS of both top 10% and bottom 10% expressed genes, that fell in a given distance interval near the TSS. For gene bodies, we performed a similar analysis by counting all the unique DSB ends mapped within the gene body of the top 10% and bottom 10% expressed genes, and normalizing the counts by gene length.

### Gene ontology analysis of top fragile genes

We identified top 10% fragile genes in three biological replicates of mouse liver tissue sections either based on the number of unique DSB ends mapped in a ±1kb interval centered on the TSS of all genes or based on the number of unique DSB ends mapped within the gene bodies. We performed GO process analysis of the fragile genes identified in all the three biological replicates, using the publicly available web-based Gorilla tool (http://bmcbioinformatics.biomedcentral.com/articles/10.1186/1471-2105-10-48).

### Identification of SpCas9, AsCpf1, LbCpf1 on- and off-target DSBs

We updated the original DSB detection pipeline previously described for analyzing Cas9-BLESS data^1,2^ to determine whether we could enhance the sensitivity of off-target detection by both BLESS and BLISS. Previously, we demonstrated that a homology search algorithm was capable of separating *bona fide* Cas9 induced DSBs from background DSBs, and performed the analysis on the top 200 DSB loci with the strongest signal after initial filtering^2^. To achieve even greater sensitivity, here we extended this homology search to the top 5,000 DSB locations identified by BLISS. To enable a direct comparison between BLESS and BLISS, we used this updated approach to re-analyze the BLESS data previously obtained with wild-type SpCas9^2^ on the same EMX and VEGFA guide targets as studied here. Briefly, a ‘Guide Homology Score’ was determined using an algorithm that searched for the best matched guide sequence within a region of the genome 50nt on either side of the center of a DSB cluster identified in BLESS/BLISS for all NGG and NAG PAM sequences in the case of SpCas9^3^, and all possible PAMs in the case of AsCpf1 and LbCpf1 for maximum sensitivity. A score based on the homology was calculated using the Pairwise^2^ module in the Biopython Python package with the following weights: a match between the sgRNA and the genomic sequence scores +3, a mismatch is –1, while an insertion or deletion between the sgRNA and genomic sequence costs –5. Thereby, an on-target sequence with the fully matched 20bp guide would have a Guide Homology Score of 60. Previously, we included the PAM match in the scoring, yielding a maximum score of 69, but in order to make the score more versatile and comparable across different PAMs, we removed the PAM dependence in the scoring. Using this guide homology score, we performed a receiver operating characteristic (ROC) curve analysis based on validated and non-validated off-targets from SpCas9-BLESS^1^ which justified our previous choice of a homology score cutoff (41 out of a max score of 60), to maximize the sensitivity and specificity of Cas9-BLISS and Cpf1-BLISS. In practical terms, this score corresponds to ≤4 mismatches or ≤2 gaps as well as combinations thereof.

